## SupplementaryData for "Differential expression of some termite neuropeptides and insulin/IGF-related hormones and their plausible functions in growth, reproduction and caste determination"

**Jan A. Veenstra**

### Table of contents

| | |
|---|---|
| SRAs used | page 2 |
| SRAs used for figure 5 | page 4 |
| Figure S1 | page 5 |
| Figure S2 | page 6 |
| Figure S3 | page 7 |
| Figure S4 | page 8 |
| Figure S5 | page 9 |
| Figure S6 | page 10 |
| Figure S7 | page 11 |
| Figure S8 | page 12 |
| Figure S9 | page 13 |
| Figure S10 | page 14 |
| Figure S11 | page 15 |
| Figure S12 | page 16 |
| Figure S13 | page 17 |
| Figure S14 | page 18 |

**SRAs used:**

*Coptotermes formosanus* : SRR2155575, SRR2155576, SRR2155577, SRR2155578, SRR10273592, SRR10273593, SRR10273594, SRR10273595, SRR10273596 and SRR10273597.
*Cryptocercus meridianus* : SRR12393556, SRR12393557, SRR12393558, SRR12393559 and SRR12393572.
*Cryptocercus punctulatus* : DRR058700, DRR058701, DRR058702 and DRR058703.
*Cryptocercus wrighti* : SRR921587.
*Cryptotermes secundus* : ERR2615943, ERR2615944, ERR2615945, ERR2615946, ERR2615947, ERR2615948, ERR2615949, ERR2615952, ERR2615950, ERR2615951, ERR2615953, ERR2615954, SRR5457739, SRR5457740, SRR5457741, SRR5457742, SRR5457743, SRR5457744, SRR5457745, SRR5457746, SRR5457747, SRR5457748, SRR5457749, SRR12381713, SRR12381714, SRR12381715, SRR12381716, SRR12381717, SRR12381718, SRR12381719, SRR12381720, SRR12381721, SRR12381722, SRR12381723, SRR12381724, SRR12381725, SRR12381726, SRR12381727, SRR12381728, SRR12381729, SRR12381730, SRR12381731, SRR12381732, SRR12381733, SRR12381734, SRR12381735, SRR12381736, SRR12381737, SRR12381738, SRR12381739, SRR12381740, SRR12381741, SRR12381742, SRR12381743, SRR12381744, SRR12381745, SRR12381746, SRR12381747, SRR12381748, SRR12381749, SRR12381750, SRR12381751, SRR12381752, SRR12381753, SRR12381754, SRR12381755, SRR12381756, SRR13221217, SRR13221218, SRR13221219, SRR13221220, SRR13221221, SRR13221222, SRR13221223, SRR13221224, SRR13221225, SRR13221226, SRR13221227, SRR13221228, SRR13221229, SRR13221230, SRR13255030, SRR13255031, SRR13255032, SRR13255033, SRR13255034, SRR13255035, SRR13255036, SRR13255037, SRR13255038, SRR13255039, SRR13255040, SRR13255041, SRR13255042, SRR13255043, SRR13255044, SRR13255045, SRR13255046, SRR13255047, SRR13255048, SRR13255049, SRR13255050, SRR13255051, SRR13255052, SRR13255053, SRR13255054, SRR13426400, SRR13426401, SRR13426402, SRR13426403, SRR13426404, SRR13426405, SRR13426406, SRR13426407, SRR13426408, SRR13426409, SRR13426410, SRR13426411, SRR15145186, SRR15145187, SRR15145188, SRR15145189, SRR15145190, SRR15145191, SRR15145192, SRR15145193, SRR15145194, SRR15145195, SRR15145196, SRR15145197, SRR15145198, SRR15145199, SRR15145200, SRR15145201, SRR15145202, SRR15145203, SRR15145204, SRR15145205, SRR15145206, SRR15145207, SRR15145208, SRR15145209, SRR15145210, SRR15145211, SRR15145212, SRR15145213, SRR15145214, SRR15145215, SRR15145216 and SRR15145217.
*Cryptotermes domesticus* : SRR2039534.
*Globitermes sulphureus* : SRR7170939, SRR9968581 and SRR12393576.
*Hodotermopsis sjostedti* : DRR084180, DRR084181, DRR084182, DRR084183, DRR084184, DRR084185 and DRR084186.
*Labritermes buttelreepeni* : SRR9968587.
*Macrotermes barneyi* : SRR3178362, SRR3182796, SRR3182798, SRR3182799, SRR3182800, SRR3182801, SRR3182802, SRR3182803, SRR3182804, SRR3182805, SRR3182806, SRR3182808, SRR3182809, SRR3182810 and SRR3182811.
*Macrotermes bellicosus* : ERR2532157, ERR2532158, ERR2532159, ERR2532160, ERR2532161, ERR2532162, ERR2532163, ERR2532164, ERR2532165, ERR2532166, ERR2532167, ERR2532168, ERR2532169, ERR2532170, ERR2532171, ERR2532172, ERR2532173, ERR2532174, ERR2532175, ERR2532176, ERR2532177, ERR2532178, ERR2532179, ERR2532180, ERR2532181, ERR2532182, ERR2532183, ERR2532184, ERR2532185, ERR2532186, ERR2532187, ERR2532188, ERR2532189, ERR2532190, ERR2532191, ERR2532192, SRR14432658, SRR14432659, SRR14432660,

SRR14432661, SRR14432662, SRR14432663, SRR14432664, SRR14432665, SRR14432666, SRR14432667, SRR14432668, SRR14432669, SRR14432670 and SRR14432671.
*Macrotermes natalensis* : SRR789255, SRR789327, SRR789336, SRR789341, SRR789345, SRR789352, SRR789356, SRR789363, SRR789369, SRR789377, SRR5457275, SRR5457276, SRR5457277, SRR5457278, SRR5457279, SRR5457280, SRR5457281, SRR5457282, SRR5457283, SRR5507526, SRR5507527, SRR13396076, SRR13396077, SRR13396078, SRR13396079, SRR13396080, SRR13396081, SRR13396082, SRR13396083, SRR13396084, SRR13396085, SRR13396086, SRR13396087, SRR13396088, SRR13396089, SRR13396090, SRR13396091, SRR13396092, SRR13396093, SRR13396094, SRR13396095, SRR13396096, SRR13396097, SRR13396098, SRR13396099 and SRR13396100.
*Mastotermes darwiniensis* : SRR921616, SRR6869961, SRR8924830, SRR8924831 and SRR12393550.
*Nasutitermes takasagoensis* : DRR162556, DRR162557, DRR162558, DRR162559, DRR162560 and DRR162561.
*Neotermes castaneus* : SRR12393532, SRR12393533, SRR12393534, SRR12393535, SRR12393536, SRR12393537, SRR12393538, SRR12393539, SRR12393540, SRR12393541, SRR12393542, SRR12393543, SRR12393544, SRR12393545, SRR12393546, SRR12393547, SRR12393548, SRR12393549, SRR12393551, SRR12393560, SRR12393562, SRR12393563, SRR12393564, SRR12393565, SRR12393566, SRR12393567, SRR12393568, SRR12393569, SRR12393570, SRR12393571 and SRR12393573.
*Prorhinotermes opinatus* : SRR12393582.
*Prorhinotermes simplex* : SRR921637, SRR13236743, SRR13236744, SRR13236745, SRR13236746, SRR13236747, SRR13236748 and SRR13236749.
*Reticulitermes aculabialis* : SRR9140410, SRR9140411, SRR9140412, SRR9140413, SRR9140414, SRR9140415, SRR9140416, SRR9140417 and, SRR9140418.
*Retitculitermes chinensis* : SRR10604052, SRR10604053, SRR10604054, SRR10604055, SRR10604056, SRR10604057, SRR10604058, SRR10604059, SRR10604060, SRR10604061, SRR10604062, SRR10604063, SRR18067184, SRR18067185, SRR18067186, SRR18067187, SRR18067188 and SRR18067189.
*Retitculitermes flavipes* : SRR5341585, SRR5341586, SRR5341587, SRR5341588, SRR5341589, SRR5341590, SRR5341591, SRR5341592 and SRR5341593.
*Retitculitermes labralis* : SRR5801942, SRR5808263, SRR8707277, SRR8707278, SRR8707279, SRR9301201, SRR9301202, SRR9301203, SRR9301204, SRR9301205, SRR9301206, SRR9301207, SRR9301208 and SRR9301209.
*Retitculitermes speratus* : DRR030795, DRR030796, DRR030797, DRR030798, DRR030799, DRR030800, DRR030801, DRR030802, DRR030803, DRR030804, DRR030805, DRR030806, DRR030807, DRR030808, DRR030809, DRR030810, DRR030811, DRR030812, DRR030813, DRR030814, DRR030815, DRR030816, DRR030817, DRR030818, DRR030819, DRR030820, DRR030821, DRR030822, DRR030823, DRR030824, DRR030825, DRR030826, DRR030827, DRR030828, DRR030829, DRR030830, DRR030831, DRR030832, DRR030833, DRR030834, DRR030835, DRR030836, DRR030837, DRR030838, DRR030839, DRR030840, DRR030841, DRR030842, DRR030843, DRR030844, DRR030845, DRR030846, DRR030847, DRR030848, DRR030849, DRR030850, DRR030851, DRR030852, DRR030853, DRR030854, DRR090831, DRR090838, DRR090840, DRR090841, DRR090842, DRR090843, DRR090844, DRR090846, DRR090847, DRR090848, DRR090852, DRR090853, DRR090854, DRR090855, DRR090856, DRR090857, DRR090858, DRR090859, DRR090860, DRR090861, DRR090862, DRR090863, DRR090864, DRR090865, DRR252502, DRR252503, DRR252504, DRR252505, DRR266547,

DRR266548, DRR266549, DRR266550, DRR266551, DRR266552, DRR266553, DRR266554, DRR266555, DRR266556, DRR266557, DRR266558, DRR266559, DRR266560, DRR266561, DRR266562, DRR332717, DRR332718, DRR332719, DRR332720, DRR332721, DRR332722, DRR332723, DRR332724, DRR332725, DRR332726, DRR332727, DRR332728, DRR332729, DRR332730, DRR332731, DRR332732, DRR332733, DRR332734, DRR332735, DRR332736, DRR332737, DRR332738, DRR332739, DRR332740, DRR332741, DRR332742, DRR332743, DRR332744, DRR332745, DRR332746, DRR332747, DRR332748, DRR332749, DRR332750, DRR332751, DRR332752, DRR332753, DRR332754, DRR332755, DRR332756, DRR332757, DRR332758, DRR332759, DRR332760, DRR332761, DRR332762, DRR332763, DRR332764, DRR332765, DRR332766, DRR332767, DRR332768, DRR332769, DRR332770, DRR332771, DRR332772, DRR332773, DRR332774, DRR332775, DRR332776, DRR332777, DRR332778, DRR332779, DRR332780, DRR332781, DRR332782, DRR332783, DRR332784, DRR332785, DRR332786, DRR332787, DRR332788, DRR357004, DRR357005, DRR357006, DRR357007, DRR357008, DRR357009, DRR357010, DRR357011, DRR357012, DRR357013, DRR357014, DRR357015, DRR357016, DRR357017, DRR357018 and DRR357019.
*Zootermopsis nevadensis* : DRR110536, DRR110537, DRR110538, DRR110539, DRR110540, DRR110541, DRR110542, DRR110543, DRR110544, DRR110545, DRR110546, DRR110547, DRR110548, DRR110549, DRR110550, DRR139981, DRR139982, DRR139983, DRR139984, DRR139985, DRR139986, DRR139987, DRR139988, DRR151559, DRR151560, DRR151561, DRR151562, DRR151563, DRR151564, DRR151565, DRR151566, DRR151567, DRR151568, DRR151569, DRR151570, SRR863596, SRR863597, SRR863598, SRR863599, SRR863601, SRR863602, SRR863603, SRR863604, SRR863605, SRR863606, SRR863612, SRR863613, SRR1167035, SRR1167037, SRR1167039, SRR1167040, SRR1167041, SRR1167042, SRR1167043, SRR1167044, SRR1167178, SRR1167247, SRR1167255, SRR1167256, SRR3139733, SRR3139734, SRR3139735, SRR3139736, SRR3139737, SRR3139738, SRR3139739, SRR3139740, SRR3139741, SRR3139742 and SRR3139743.

**SRAs used for figure 5:**
ERR2615943, ERR2615944, ERR2615945, ERR2615946, SRR5457742, SRR5457743, SRR5457744, SRR5457745, SRR12381714, SRR12381716, SRR12381718, SRR12381721, SRR12381723, SRR12381725, SRR12381727, SRR12381728, SRR12381730, SRR12381732, SRR12381734, SRR12381736, SRR12381738, SRR12381740, SRR12381743, SRR12381744, SRR12381745, SRR12381746, SRR12381747, SRR12381748, SRR12381755 and SRR12381756.

```
Blattella      MKNFQAQVIV-ITAAILLHQCAGRPEYEDCNRKIRRQILESCSSEKGKRSAEYDTPSLPM
Periplaneta    MKPHFALVIVVVV-STLFQAGIKGEKYENCSKKLRQLILDSCNEPKNKRSAVFERHGFNM
Cryptocercus   MKSILILVIIVAT-SSLLHESFGSSDYKYCSKKMRQLILDSCSEPKGKRSAILADYNLNK
Zootermopsis   MKSFMALVIAVVTSAVVLQEGVRSSDYENCSKKMRQLILDSCAEPKGKRNAFLADFNWNI
Hodotermopsis  MKILVALVIIVVTSAFVLQEGVGISDYENCSKKMRQLILDSCAEPKGKRNAFLADYNWNV
Cryptotermes   MKELLVLIIVVMTSAHLLQESVANSHYENCSKKMRQLILDSCAEPKDKRNAILAENKDLF
Macrotermes    --------------SILLPERVANSDYENCSKKLRQLVHNSCAESLDKRRAILVQD---I
               1........10........20........30........40........50.........

Blattella      HD-----EPLQNAPS-SALLGRILGVPSQWTADDVAVNNANRQVKRSPETIRQLMIDCCL
Periplaneta    HSPHLPQDRSQQVTSS-VLLGKILGVPSQWTEEELSSHQINKQFRRNNQSVRNLIIECCV
Cryptocercus   FYPHSVHIPVRHIASP-AMLGKILGVPSHWTEDLVSLDDSSQQYKRTLPEENDLIVMCCA
Zootermopsis   HP---PRKAAQHTVSP-ALLGKVLGVPPHWMEGVVSLDDSKQPQRRNLQAVHNLIVECCV
Hodotermopsis  PP---PHTPARHTASP-ALLGKLLGVPPHWTEGLVSLDDSQ-QQRRNLQAVHNLIVECCV
Cryptotermes   -T---RHRPAQQSASPSLLAGKVLGVPSCWTEDLAIFDDSNIQHRRNLPAVHNLIVECCV
Macrotermes    FYTDAPNKPARQTPSP--SAGKILCVPSHWTDDLASFDDTNKQHRRKLPEVQN*IVECCV
               61.......70........80........90........100.......110........

Blattella      ANCSPDRFLGMC*----------
Periplaneta    DGCTPNQIMGLCD*---------
Cryptocercus   NGCDPSKIVGLCN*---------
Zootermopsis   DGCTPYQIVGLCN*---------
Hodotermopsis  DGCTPNQIVGLCN*---------
Cryptotermes   DGCTPSQILGLCN*---------
Macrotermes    DGCSPRQVLGLCNRTHRFSFKA*
               121......130.......140.
```

**Figure S1.** Sequence comparison of predicted gonadulin sequences from three cockroach species, *Blattella germanica*, *Periplaneta americana* and *Cryptocercus wrighti* and four termite species, *Zootermopis nevadensis, Hodotermopsis sjostedti*, *Cryptotermes secundus* and *Macrotermes natalensis.* Note that the *Macrotermes* sequence is derived from a pseudogene as it has an inframe stop codon and appears to lack the first part of the signal peptide. Also note the unusual seventh cysteine in the *Cryptotermes* sequence. Identical amino acid residues are highlighted in black, conservative substitutions are in grey and the cysteine residues are highlighted in red. An inframe stop codon in the *Macrotermes* sequence is highlighted in yellow.

```
Blattella       MLKCGIVTALVLVTTMVSGAPTIRMQMCGSQLANTLAQICSAYGYHDPFSQTRRVNSPSS
Periplaneta     ----MS-LLFLVLLASATASPLGRMQLCGSELANKLAEICAVYGYNDPFRHAHY-ED-ST
Zootermopsis    -MPLLSVVRIFLLTMTVSASPLGRLQLCGSELANKLAEICSIQGYNDPFRHSMYYEV-SP
Hodotermopsis   -MSLLSVVRIFLLTMTVTASPLRRMQLCGSELANKLAEICSIQGYNDPFRQSMYYEI-SP
Cryptotermes    MPLALSLVRVFLLTMTVAASPLGRLQLCGSELANKLAEICSVQGYNDPFRNSMYYEI-SP
Coptotermes     MSLHLCVVRMLLLTMTVAASPLGMLQLCGSALANKLAEICAVQGYNDPFKHSMYYDM-SP
Prorhinotermes  MSLHLCVVRILLVTMTVASSPLGKLQLCGSDLANKLAEICAAQGYNDPFKHSMYYEM-SP
Reticulitermes  MSLHLCVVRMLLLTMTVAASPLGMLQLCGSALANKLAEICAAQGYNDPFKHSMYYEM-SP
Macrotermes     MSPHLSVVRILLLTVTVAASPLGRLQLCGSALANKLAEICSVQGYNDPFRHSMYYEM-SS
Labritermes     MSPHLSVARVLLLTVAVAASPLGRLQLCGSDLANKLAEICAAQGYNDPFRHSMYYEL-SP
Nasutitermes    ----------------------------------------VQGYNDPFRHNMYYEM-SP
                1.......10.......20.......30.......40.......50.........

Blattella       GVNTTPNRLRVRRGVADECCKTGCTLDTMEQYCSAPLTPAQRARFLQQYQSNALNRILQE
Periplaneta     YDSITPTRTRVKRGVADECCKTGCSQETLEQYCNPPLKSPDRSKVLKISEDHSINHIPQD
Zootermopsis    FEGGVSSTTRSKRGVADECCKTGCSWDTLEQYCNPPLSTPDRPHNLQETEDHSINHIAQV
Hodotermopsis   FGGVVPLTTRSKRGVADECCKTGCSWDTLEQYCNPPLSTPDRPHNLQETEDHSTNHISQD
Cryptotermes    LDNGVPLMTRSKRGVADECCKTGCSWDTLELYCNPPLPTAQRPHNLQEAEDHSMNHIPLD
Coptotermes     FENGIPITTRSKRGVADECCKKGCSWETLEQYCNPPLPTPSRTDNLQATEEHSMNRIPHD
Prorhinotermes  FENGIPMTTRSKRGVADECCKKGCSWDTLEQYCNPPLRTPNRTDNLQATEEHSXX-----
Reticulitermes  LDNGIPVTTRSKRGVADECCKKGCSWDTLEQYCNPPLPAPNRTDNLQATEEHSMNRIPQD
Macrotermes     LENGIPMTTRSKRGVADECCKKGCSWDALEQYCNPPNPTRNRTDNLQATEEHSMNRIPQD
Labritermes     LENSIPMTTRGKRGVADECCKKGCSWDTLEQYCNPPIPTPNRTDNLQSTEEQSINRIPHD
Nasutitermes    LENGIPMTTRAKRGVADECCKKGCSWNTLEQYCNPPIPSSNRTDILQETEEQSMNRIPHD
                61.......70.......80.......90.......100.......110........

Blattella       VPSGSSLN----AGSKISKDPKNDLSSKVRRQQDKKGQRGNNRCRCRRRRRRGKGDSEEI
Periplaneta     DSAASAVLTSEVRRSSIAREEKNDLVSKVRGHHDKKGRRANNRCRCRRRRRRGKGDLEEL
Zootermopsis    DSEGSAG-----QPTTITREQKNDLVSKVRSHHEKKGRRGNNRCRCRRRRRRGKGDIEDP
Hodotermopsis   VSEGSAG-----LPPATTREQKNDVVSKVRSHHDRKGRRGNNRCRCRRRRRRGKGDIEDP
Cryptotermes    DSTGSEA-----RPPAKSRAQKNDLVSKVRSHHDKKGRRGNNRCRCRRRRRRGKGDMEDA
Coptotermes     DSAGSAG-----RLPLRARDQKSDLISKVRSHHEKKGRRGNNRCRCRRRRRRGKGDIEDP
Prorhinotermes  ---------------------------XXRSHHEKKGRRGNNRCRCRRRRRRGKGDTEDP
Reticulitermes  DSAGSAA-----RPPIMTRDQKSDLISKVRSHHEKRGRRGNNRCRCRRRRRRGKGDIEDP
Macrotermes     DSAGTAG-----RPSIMTRDQKSDLISKVRSHHEKKGRRGNNRCRCRRRRRRGKGDIEDP
Labritermes     DSASSAA-----RPQIMTRDQKSDLISKVRSHNEKKGRRANNRCRCRRRRRRGKGDIEDP
Nasutitermes    DSASSAA-----RAPITTRDQKSDLISKVRSHHEKKGRRGNNRCRCRRRRRRGKG-----
                121.....130.......140.......150.......160.......170........

Blattella       ERQQNHIAPVIGTINPSYFGVPVFLSPRLKKQETQQDRHRK*-
Periplaneta     ERHLNKIAPVIGTINPSYLGIPVILPPRIRKEEMSLQDYGTK*
Zootermopsis    EILENKIAPVIGTINPAYFANPVILPPRIRKEEMALQDTK*--
Hodotermopsis   EIFENKIAPVIGTINPSYFANPVILPSRIRKEEMSLQDTK*--
Cryptotermes    EILENKIAPVIGTINPSYFANPVILPPRVRKEEMSLQDRK*--
Coptotermes     EVLENKIAPVIGTINPSYFANPVILSQRIRKEETSLQETK*--
Prorhinotermes  EVLENKIAPVIGTINPSYFGNPVILPPRIRKEEMSLQEKK*--
Reticulitermes  EVLENKIAPVIGTINPSYFANPVILPPRIRKEEMSLQETK*--
Macrotermes     EVLENKIAPVIGTINPSYFANPVILPPRIRKEEMSLQETQ*--
Labritermes     EVLENKIAPVIGTINPSYFANPVILPPRIRKEGMSLQETQ*--
Nasutitermes    -------------------------------------------
                181.....190.......200.......210.......220.
```

**Figure S2.** Sequence alignment of the predicted long IGFs from two cockroaches, *Blattella germanica* and *Periplaneta americana*, and nine termite species, *Zootermopis nevadensis, Hodotermopsis sjostedti, Cryptotermes secundus, Coptotermes formosanus, Prorhinotermes simplex, Reticulitermes speratus, Macrotermes natalensis, Labritermes buttelreepeni* and *Nasutitermes takasagoensis.* Note that the *Prorhinotermes* and *Nasutitermes* sequences are incomplete due to lack of data. Highlighting of residues as in figure S1.

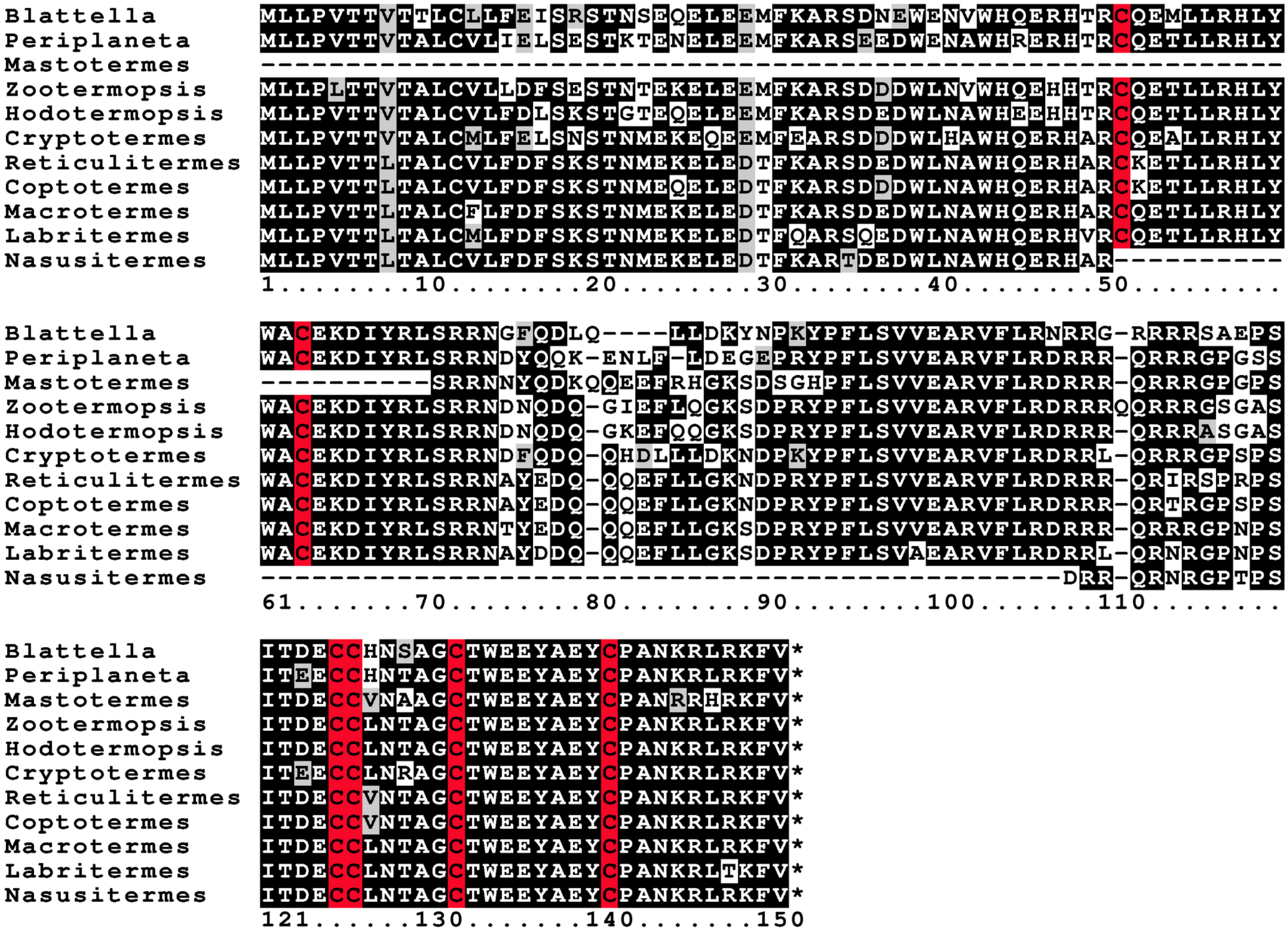


**Figure S3.** Sequence alignment of the predicted dilp7 orthologs from two cockroaches, *Blattella germanica* and *Periplaneta americana*, and nine termite species, *Mastotermes darwiniensis, Zootermopis nevadensis, Hodotermopsis sjostedti, Cryptotermes secundus, Coptotermes formosanus, Reticulitermes speratus, Macrotermes natalensis, Labritermes buttelreepeni* and *Nasutitermes takasagoensis.* Note that the *Mastotermes* and *Nasutitermes* sequences are incomplete due to lack of data. Highlighting of residues as in figure S1.

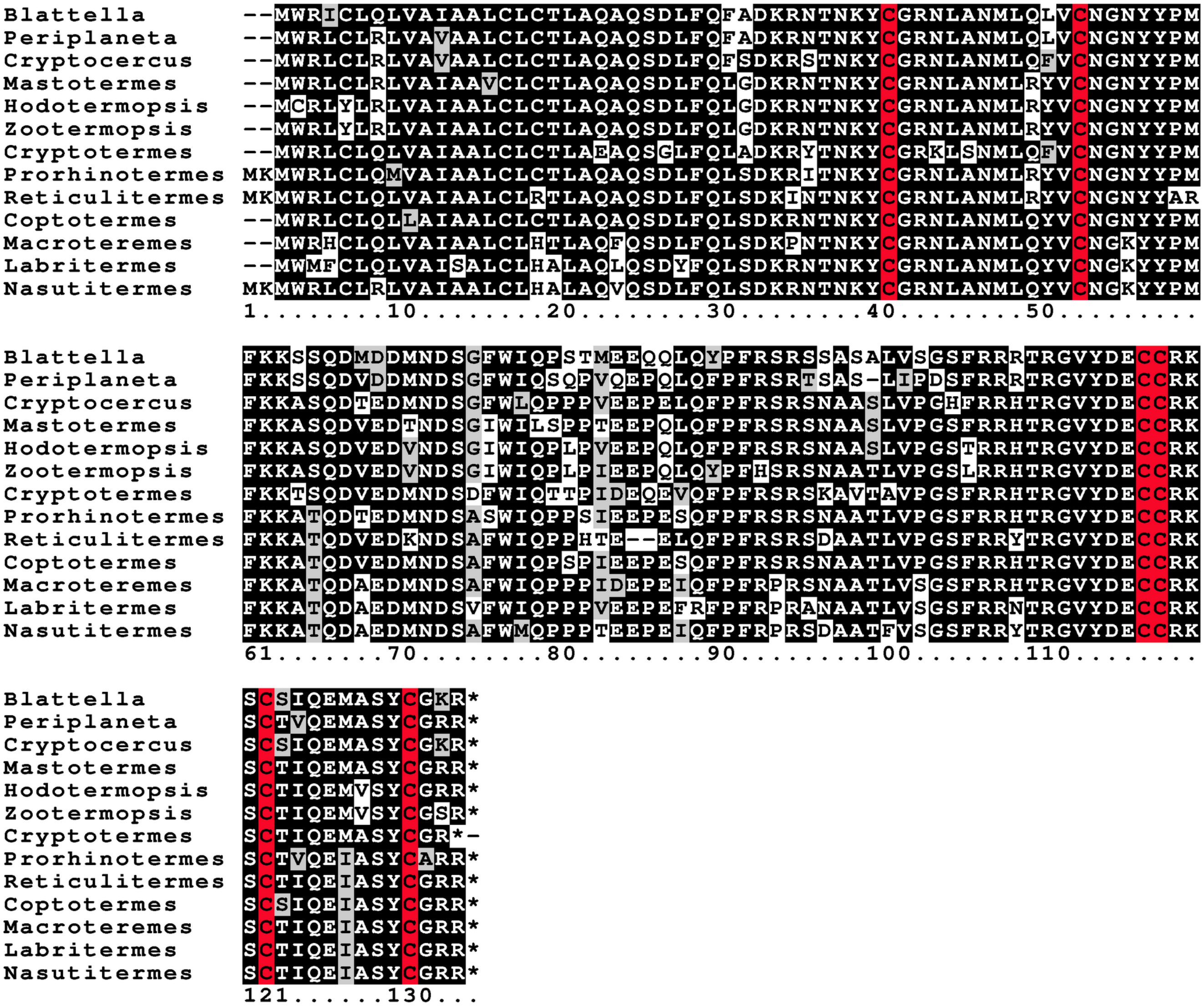


**Figure S4.** Sequence alignment of the predicted atripins from three cockroach species, *Blattella germanica*, *Periplaneta americana* and *Cryptocercus wrighti* and ten termite species, *Mastotermes darwiniensis*, *Zootermopis nevadensis*, *Hodotermopsis sjostedti*, *Cryptotermes secundus*, *Prorhinotermes simplex*, *Coptotermes formosanus*, *Reticulitermes speratus*, *Macrotermes natalensis*, *Labritermes buttelreepeni* and *Nasutitermes takasagoensis*. Highlighting of residues as in figure S1.

```
Periplaneta-3    ----MWRV-VMVAACLCVLCESQSDLFQPQDKR-GSRRYCGPTLVSTLHIICNGTYYTNI
Periplaneta-4    ----MLKVAVLMAACLCVLCESQSDML---EKRDSARRYCGRNLADVMHIVCNGIYNGNP
Cryptocercus     MWRSCLRALVVVALCLCSMSESQSDVFQFQDKRAEAKRYCGNNLVAALQFLCDGRYYSNF
Mastotermes      MWRVCFRVMVVVAMCLCSLPQSQSDVLQFPDKRQEAKRYCGDNLVRILQFLCDGIYYSNT
Hodotermopsis    MWRACFRVMVVVAMCLCSLAQSQSDIFQFPDKRPETKRYCGSNLVDILQLLCNGKYYSNT
Zootermopsis     MWRACFRIVVVVALCLCSLAQSQSDIFPFPDKRPETKRYCGSNLVDILQLVCNGKYYSNI
Cryptotermes     MWRACFTVMVVVAMCLCSLADSQSDLYQFPSKRPDSKRYCGSNLVKILQFLCDGYYNANI
Prorhinotermes   MWKACFRMIVFVAMCLCPLAESQSDIFQFPDKRLEARRYCGNNLVDKLQLLCNGYYYSNI
Reticulitermes   MWRVCFRMMVFAAMCLCSLAESQSDIFQFPDKRLETKRYCGNNLVDILQFLCNGYYYSNI
Coptotermes      MWRVCFRMTVFAAMCLCSLAESQSDIFQFPDKRPETKRYCGNNLVDILQLLCNGYYYSNI
Macroteremes     MWRACFRMMVFVAMCLCSLAESQSDIFQFPDKRLESKRYCGSNLVDMLQFLCNGYYYSNI
Labritermes      MWRVCFRTTVFVAVCLCSLAESQSDIFQFPDKRLENKRYCGNNLVDILQFLCNGYYYSNI
Nasutitermes     MWRSCFRMMVLVAMCLCSLAGSQSDIFQFPDKRLENKRYCGNNLVDILQFVCNGYYYSNI
                 1.......10........20........30........40........50.........

Periplaneta-3    RPL----------SRAQKKAWPDSDDDVWMELQLPENAERYPFRSRASAVAFPRRVFKR-
Periplaneta-4    R------------AATQKKSWPDADDDLW----PVEEQADFPFRSRASA----HRILKRQ
Cryptocercus     NTN---NSNNFNQHAGQKKSVSEMDDDYWLKLQPVEEHLKFPFRSR----TFAHKIFKR-
Mastotermes      NLNNNN-----NQRVGRKKSVPEVDEDFWLQLQPVEEQMKFPFLSRSSASTFAHRIFKR-
Hodotermopsis    NSNN--SNNNYNPHVGRKKSMPEVEEDFWQQFQPVEEQMKFPFRSRSSSSAFAHRIFKR-
Zootermopsis     NN-----SNNYSPHVGRKKSMPEADEDFWQQLQPVEEQMKFPFRSRSSVSTFAHRIFKR-
Cryptotermes     S------PNNYGTRVGRKKSIPEADDDLWLQLQPAEEEIKFPFRPRSSASIFSHRMFKR-
Prorhinotermes   NANNNNNNNNYSPHAGRKKSMPEVDEDLWHQLQPVEEHMTFPFRSRSSASTFLHRIFKR-
Reticulitermes   NSN---NSNNYSPHTGRKKSMPEVDEDFWHQLQPLEEHMKFPFRSRSSASTFLHRIFKR-
Coptotermes      NSN---TSNNYSPHAGRKKSMPEVDEDFWHELQPAEEHMKFPFRSRSSASTFLHRIFKR-
Macroteremes     NSN---NSNNYSPHAGRKKSMPEVDEDLWHELEPVAEHIKFPFRSRSSASTFLHRIFKR-
Labritermes      NSS---NSNNYSPHAGRKKSVPEADEDLWHELEPVEEHMKFPFRSRSSASTFLHRIFKR-
Nasutitermes     NSN---NSYNYSPHAGRKKSVPEVDEDLWREIEPVEEHLKFPFRSRSSASTFLHRIFKR-
                 61.......70........80........90........100.......110........

Periplaneta-3    QSWGVADECCVRKGCDYNELSSYCAP*----
Periplaneta-4    SPIGIAYECCINKGCTIHELRTYCGR*----
Cryptocercus     HPGGVAYECCIIKGCTINELMSYCMPSTEQ*
Mastotermes      HPVGVAYECCISKGCTVQELSSYCGRS*---
Hodotermopsis    HTVGVAYECCINKGCTMHELRSYCAP*----
Zootermopsis     HTVGVAYECCINKGCTVYELRSYCAP*----
Cryptotermes     HSVGVAYECCINKGCTVDELKSYCGR*----
Prorhinotermes   HAVGVAYECCISKGCTIHELRGYCAP*----
Reticulitermes   HAVGVAYECCISKGCTIHELRGYCGP*----
Coptotermes      HAVGVAYECCISKGCTIHELRSYCGP*----
Macroteremes     HAVGVAYECCISKGCTIPELRSYCGS*----
Labritermes      HAVGVAYECCISKGCTIHELRSYCAP*----
Nasutitermes     HAVGVAYECCISKGCTIHELRSYCGP*----
                 121......130.......140.........
```

**Figure S5.** Sequence alignment of the predicted birpins from two cockroach species, *Periplaneta americana* and *Cryptocercus wrighti* and ten termite species, *Mastotermes darwiniensis, Zootermopis nevadensis, Hodotermopsis sjostedti, Cryptotermes secundus, Prorhinotermes simplex, Coptotermes formosanus, Reticulitermes speratus, Macrotermes natalensis, Labritermes buttelreepeni* and *Nasutitermes takasagoensis.* Note that these sequences are not as well conserved as the atirpin sequences (Fig. S4). Highlighting of residues as in figure S1.

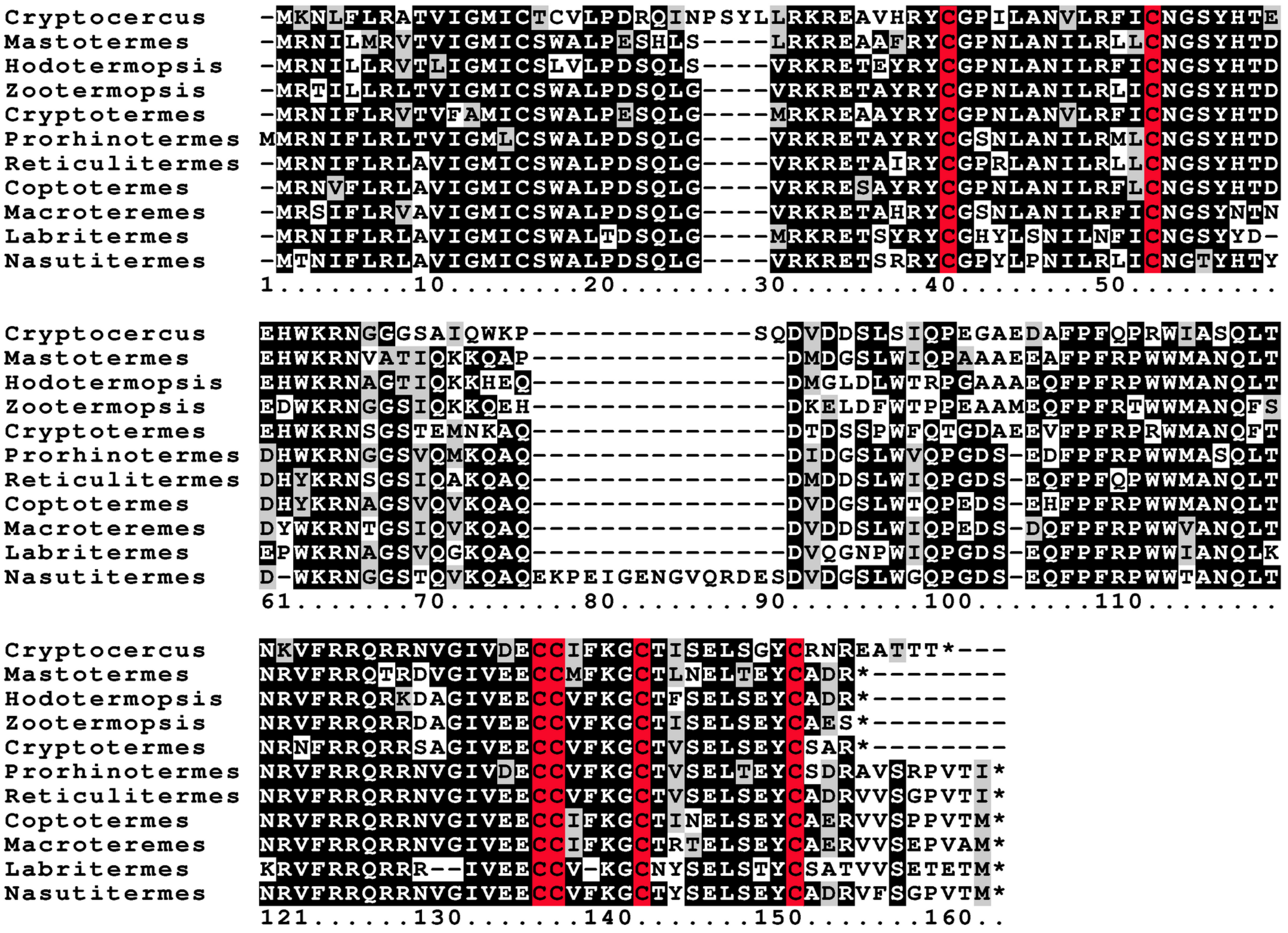


**Figure S6.** Sequence alignment of the predicted cirpins from one cockroach species, *Cryptocercus wrighti* and ten termite species, *Mastotermes darwiniensis, Zootermopis nevadensis, Hodotermopsis sjostedti, Cryptotermes secundus, Prorhinotermes simplex, Coptotermes formosanus, Reticulitermes speratus, Macrotermes natalensis, Labritermes buttelreepeni* and *Nasutitermes takasagoensis.* Note that these sequences are not as well conserved as the atirpin sequences (Fig. S4). Highlighting of residues as in figure S1.

```
Periplaneta-2      MWKLCLLVM----VTCACALPEN-------LNKRDSSHRYCGRNLVNILSFVCSGRGYNG
Cryptocercus       MWKLCLRVA-LIGLICVCALPENPSTMLQLVSKRETTHRYCGKILVNTLRFLCRSQ-YNG
Mastotermes        MRRLCLHVM-LIGVICVCALPENPSAVLQLVRKRDATHRYCGSNLVTTLRFLCRSQ-YNG
Hodotermopsis      MWRLCSRVMMLIGIICVCALPDHPSTAAKIVQKRQTTHRYCGKNLVDVLKFLCRGQ-YNG
Zootermopsis       MWRLCLRAM-LISMICVCALTDNPSTTPKFVEKR-EAHRYCGRNLANILKFLCQGH-YNG
Cryptotermes       MWRLFLRVM-LIGVICACALPENPSFI-KSVRKRDTSQRYCGKNLVNILRFLCRSQ-YNG
Prorhinotermes     MWRLCLPVM-LIGAICVCSLPGDPSSMSKLVTKREAEQRYCGTKLAAILRYVCNGT-YFA
Reticulitermes-1   MWRLSLRVM-LIAVICVCVLSGIPSSMSKYVNKREKEQRYCGSKLADKLREVCNGI-YES
Reticulitermes-2   MARLWLRVM-LIAVIGVCAVFEKPSSMSKYVKKRETEERYCGSKLADKLREVC-------
Reticulitermes-3   MWRLCLRVM-LIGVICVCALPENPSLMSKFVKKRETQQRFCGSKLVAILRLLCQGK-YHG
Coptotermes-1      MWRLCLSVV-LIGVICVCALPENPSLMSKLVQKRETQQRLCGNNLVAILRLLCQSK-YHG
Coptotermes-2      MWRLWLRVM-LITAICACAVFEKPSSMSKSVKKRQTEERYCGSKLADKLMEVCNGT-YEG
Coptotermes-3      MWTLSLRVM-LTAVICVRVLSGIPSSMSRFVQKREIEQRYCGSKLYYKLREVCNST-YQG
Coptotermes-4      MWRQWLRVM-LITAICACAVFEKPSSMSKFVKKRQTEV*YCGSKLADKLMEVCKST-YEG
Macroteremes-1     MWRLCLRVM-LIAVICVCALPEKPSVMSKLVRKRETQQRYCGSRLVNILKFLCQSK-YYG
Macroteremes-2     MWRLCLRVM-LIAVICVCALPDKLSVMSKFLKKRETEQRYCGSRLVDKLKEVCNST-YHS
Labritermes-1      MWRLCLRVM-LVAVICVCALPAKPSVMSKLVRKRETQQRYCGSRLVHILKFLCQSK-YYG
Labritermes-2      MWKLFLRVM-LVAVICVCALPAKPSVMSKLVRKRETQQRYCGSRLVHILKFLCQSK-YYG
                   1........10........20........30........40........50.........

Periplaneta-2      VEEMEERKRN--------------------------SHALPDADDAFWWLQLS---DEEL
Cryptocercus       IEDETKRNADPIQK---------------------KSLQ--DTDEYLWIQPQAVDERQEI
Mastotermes        IDDERKRNAEPIVK---------------------KSLQVSDPDNSLWTQLQTVDELPET
Hodotermopsis      MEYERKKRDV-YYS---------------------EEDPFEDRDDDFWSQSEDLSGFPNL
Zootermopsis       IDYDRKRRDVDFYG---------------------EPKLYGDHDYDFWSQAEDVDDFADL
Cryptotermes       IEEERKRSEETSQR---------------------KVAR--DLEYVQWSRPKPVE-EPEI
Prorhinotermes     PEAKRKRDT-----------------------GFFENMQEEDSDTDLWMESQDVEEFQDA
Reticulitermes-1   MDKKMK----------------------------------RDLDTLQWIQSHAEEVVSNT
Reticulitermes-2   ----------------------------------------SDLDTVLWFQSQAEKVVSNT
Reticulitermes-3   MFERRKRDT-----------------------DFFLWRSPQDLDTLPWIQSHAEEVVSNT
Coptotermes-1      MFERRKRDT-----------------------HMFHSRSPQDLDTLQWIQSQAEEEVPKT
Coptotermes-2      PNNTDQGVNNTDLGVNNTDKGVNNTDLGVNTDQGVNTPRKRDLRTVLWFQSQAEKEVRNT
Coptotermes-3      MENKHK----------------------------------RDIHTVLWFQSRGEKEVPNT
Coptotermes-4      PNNTDQ------------------------------------------------------
Macroteremes-1     MEERRRRDT-----------------------EFFHTRSPQDMGTVLWNQSQAEEQVPNT
Macroteremes-2     MEMK-----------------------------------KRDMGTVLWNQSQVEEQVPNT
Labritermes-1      MEERRKRDT-----------------------EFFHRRSPQDMGTVLWNQSPAEEQVPNP
Labritermes-2      MEERRKRDT-----------------------EFFHRRSPQDLDNVLWNQTQTEENVPNP
                   61.......70........80........90........100.......110........

Periplaneta-2      QFPFRSRAAANTLSNREFRRQ-TRNDRGIVEECCIVKGCTINELADYCLSR*--------
Cryptocercus       QFPFRPRSIANTFANRIFRRQ-ERSDGGIVDECCIYKGCTLNELTAYCSNG*--------
Mastotermes        EFPFRSRSIANTFANRVFRRR-ERSDGGIVDECCIYKGCTVNELTEYCSSR*--------
Hodotermopsis      QYPFRSRPAANTLAN-MFRRRQK-RGDGIVEECCVSKGCTVNELRDYCSPR*--------
Zootermopsis       HYPFRPRSVSSVLLNKLFRRRPKRNNGGIVEECCIYKGCTLSELTEYCSPVQ*-------
Cryptotermes       RFPFRSRAAANT-----FRRQ-KRNDGGIVDECCIYKGCTVNELTDYCSSRR*-------
Prorhinotermes     KFPFNSHSEASTFKNRRFRRE-DGNYTGIVDECCVYKGCTLTEVAEYCAPQQPDASTTNS
Reticulitermes-1   KFPFRPRSIANKFKKRSLRRQ-KRTSGGVADECCINKGCTIGELREYCAER*--------
Reticulitermes-2   KFPFQPRSVVNKFKKRSLRRQ-KRDESGVAYECCINKGCTIGELRDYCAER*--------
Reticulitermes-3   KFPFRPRSIANKFKNRSLRRQ-KRTSGGVADECCINKGCTIGELREYCAER*--------
Coptotermes-1      KFPFRSRSVANTFKNRSLRRL-KRNSGGVADECCINKGCTLGELREYCAER*--------
Coptotermes-2      KFPFYPRSAANKFKKRSFRRQ-KR-EGGVADECCINKGCTVTELRQYCAER*--------
Coptotermes-3      KFPFRPRSVANKFS-RSLRRQ-KRNSRGVADECCINKGCTLRELREYCAQR*--------
Coptotermes-4      ---------ANKLKKRSLRRQ-KR-EGGVADECCINKSCTVTELRQYCAER*--------
Macroteremes-1     MYPFRPRSVANAFTNRSLRRQ-RRTPRGVADECCIYKGCTVSELRAYCAER*--------
Macroteremes-2     KYPFHSRSVANTFTNRSLRRQ-KRNDGDVANECCIYKGCTVSELRAYCGER*--------
Labritermes-1      KYPFRPRSLANMFKNRSLRRE-KRNDGGVADECCIKKGCIISELRAYCAER*--------
Labritermes-2      KYPFRPRSLANMFKNRSLRRQ-KRTEGGVADECCIYKGCTYSELRAYCAER*--------
                   121......130.......140.......150.......160.......170........
```

**Figure S7.** Sequence alignment of the predicted brovirpins from two cockroach species, *Periplaneta americana* and *Cryptocercus wrighti* and ten termite species, *Mastotermes darwiniensis, Zootermopis nevadensis, Hodotermopsis sjostedti, Cryptotermes secundus, Prorhinotermes simplex, Coptotermes formosanus, Reticulitermes speratus, Macrotermes natalensis, Labritermes buttelreepeni* and *Nasutitermes takasagoensis.* Note that the coding sequence for brovipirn 4 from *Coptotermes* contains as stop codon (highlighted in yellow) and is thus unlikely to be functional and that some sequences are incomplete due to lack of data. Other highlighting of residues as in figure S1.

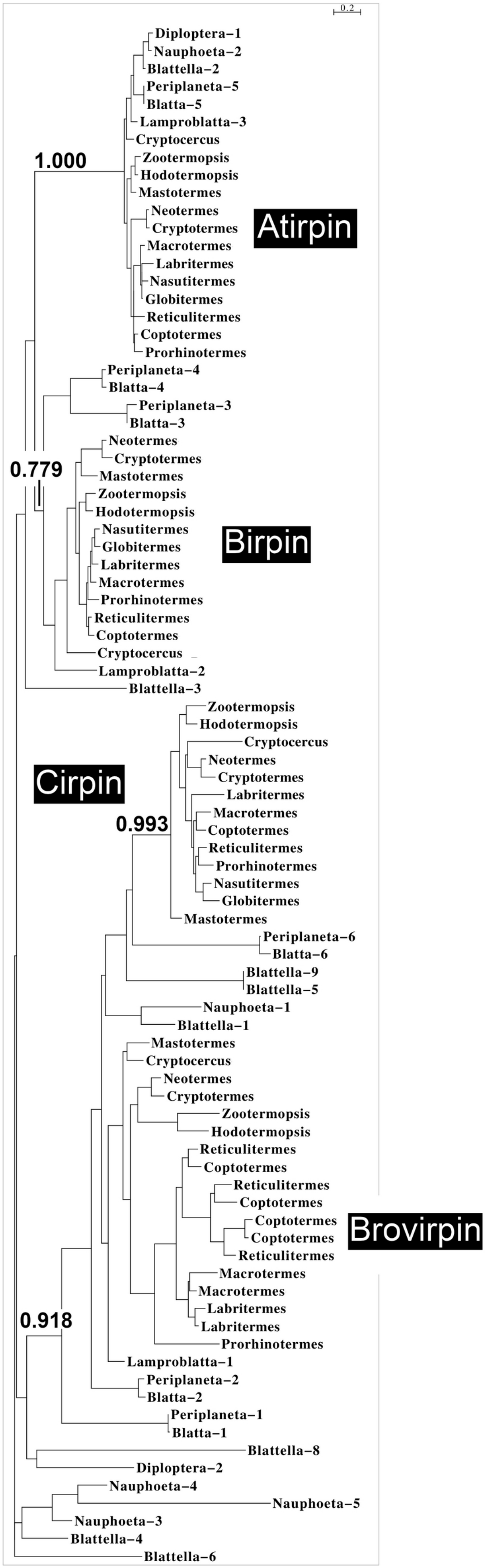


**Figure S8.** Sequence similarity tree of Blattodea sirps.

```
M.nat-aa                    ValAlaAspLeuLeuMetAlaIleTyrLeuLeuValThrGly
M.nat-nt  gtgataataatgctgttagttgcagacctattaatggcaatctatctactggtgactgga
C.sec-nt  ataatattggtttccgcagtttcagatttgttaatggcgttttatctactggtgattggg
C.sec-aa                    ValSerAspLeuLeuMetAlaPheTyrLeuLeuValIleGly

M.nat-aa  ThrGlnGlyCys***PheArgGlyHisTyrHisArgAspAlaHisSerTrpIleSerSer
M.nat-nt  actcagggttgttaattccg-ggacattatcacagagaggcacacagctggatatcgtct
C.sec-nt  atccaggactgtcagttccgaggcaactatcacaaggaggctcacaagtggatgtcatct
C.sec-aa  IleGlnAspCysGlnPheArgGlyAsnTyrHisLysAspAlaHisLysTrpMetSerSer

M.nat-aa  ***ValCysThrLeuIleGlyMetValAlaIleThrSerSerAsp
M.nat-nt  tgagtatgcacgctaattggcatggtcgcgatcacgtcatcagaagtgaatatcaaatac
C.sec-nt  tggggatgcacgctcatcggtatggtcgcaatgacgtcatcagaagtgagtctcagcctc
C.sec-aa  TrpGlyCysThrLeuIleGlyMetValAlaMetThrSerSerAsp
```

**Figure S9.** The only remaining "coding" exon of LGR3 from *Macrotermes natalensis*. Nucleotide and conceptual translated amino acid sequences are compared with those from the orthologous exon from *Cryptotermes secundus,* in which LGR3 is presumed to be functional. Highlighted in black is the predicted amino acid sequence encoded by the *S. secundus* exon as well as those amino acid residues that are converved between the two species. Although some amino acid substitutions are conservative, there are also two in-frame stop codons as well as a single nucleotide deletion (all highlighted in yellow). Hence this can not be part of a functional protein. *M.nat-aa* and *M.nat-nt M.natalensis* amino acid and nucleotide sequences respectively, *C.sec-aa* and *C.sec-nt*, the same for *C. secundus*.

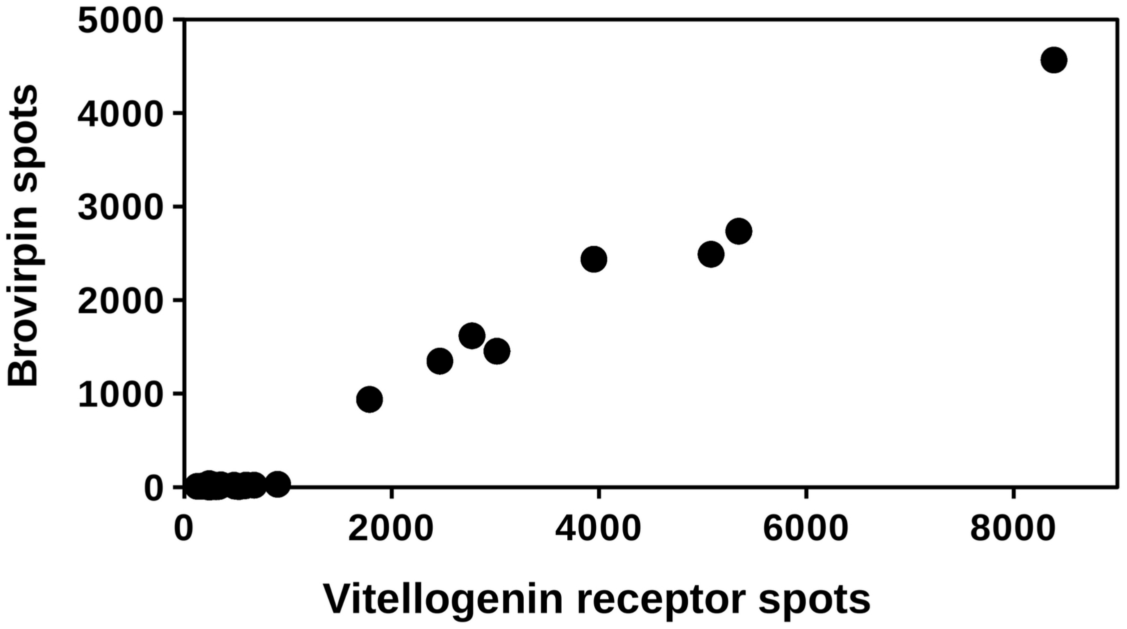


**Figure S10.** Correlation between the numbers of reads for vitellogenin receptor and brovirpin in transcriptome SRAs from *C. secundus* queens. Note that only those SRAs that have large numbers of brovirpin reads also have large numbers of vitellogenin receptor reads. The SRAs used here are ERR2615943, ERR2615944, ERR2615945, ERR2615946, SRR5457742, SRR5457743, SRR5457744, SRR5457745, SRR12381714, SRR12381716, SRR12381718, SRR12381721, SRR12381723, SRR12381725, SRR12381727, SRR12381728, SRR12381730, SRR12381732, SRR12381734, SRR12381736, SRR12381738, SRR12381740, SRR12381743, SRR12381744, SRR12381745, SRR12381746, SRR12381747, SRR12381748, SRR12381755 and SRR12381756.

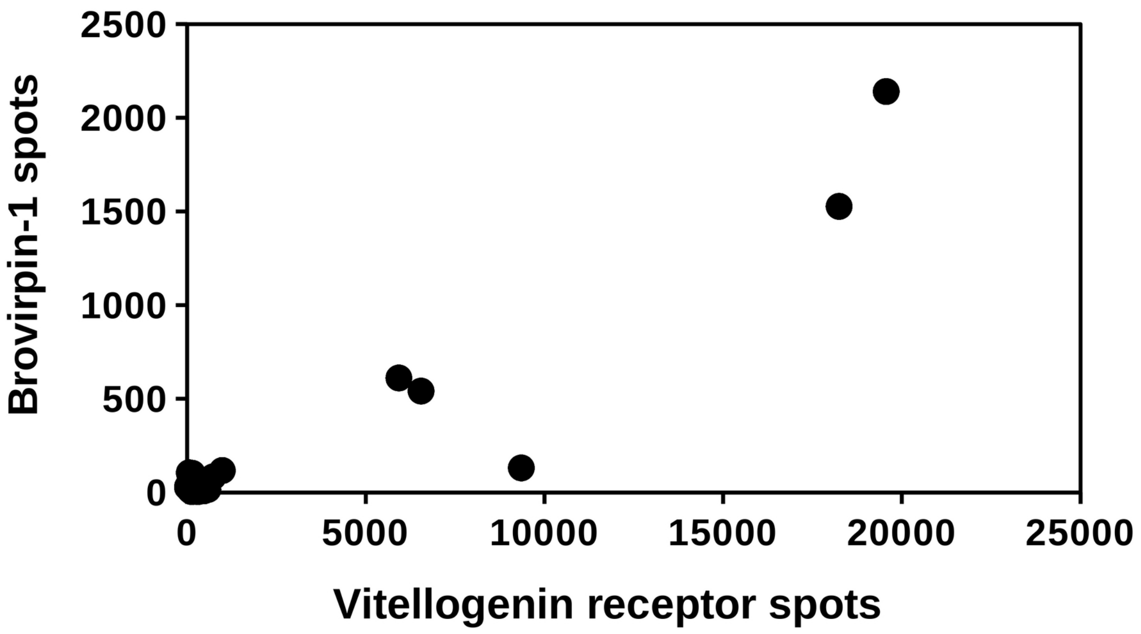


**Figure S11.** Correlation between the numbers of reads for vitellogenin receptor and brovirpin in transcriptome SRAs from *M. natalensis*. Note that only those SRAs that have large numbers of brovirpin reads also have large numbers of vitellogenin receptor reads. The SRAs used here are : SRR13396076, SRR13396077, SRR13396078, SRR13396079, SRR13396080, SRR13396081, SRR13396082, SRR13396083, SRR13396084, SRR13396085, SRR13396086, SRR13396087, SRR13396088, SRR13396089, SRR13396090, SRR13396091, SRR13396092, SRR13396093, SRR13396094, SRR13396095, SRR13396096, SRR13396097, SRR13396098, SRR13396099 and SRR13396100.

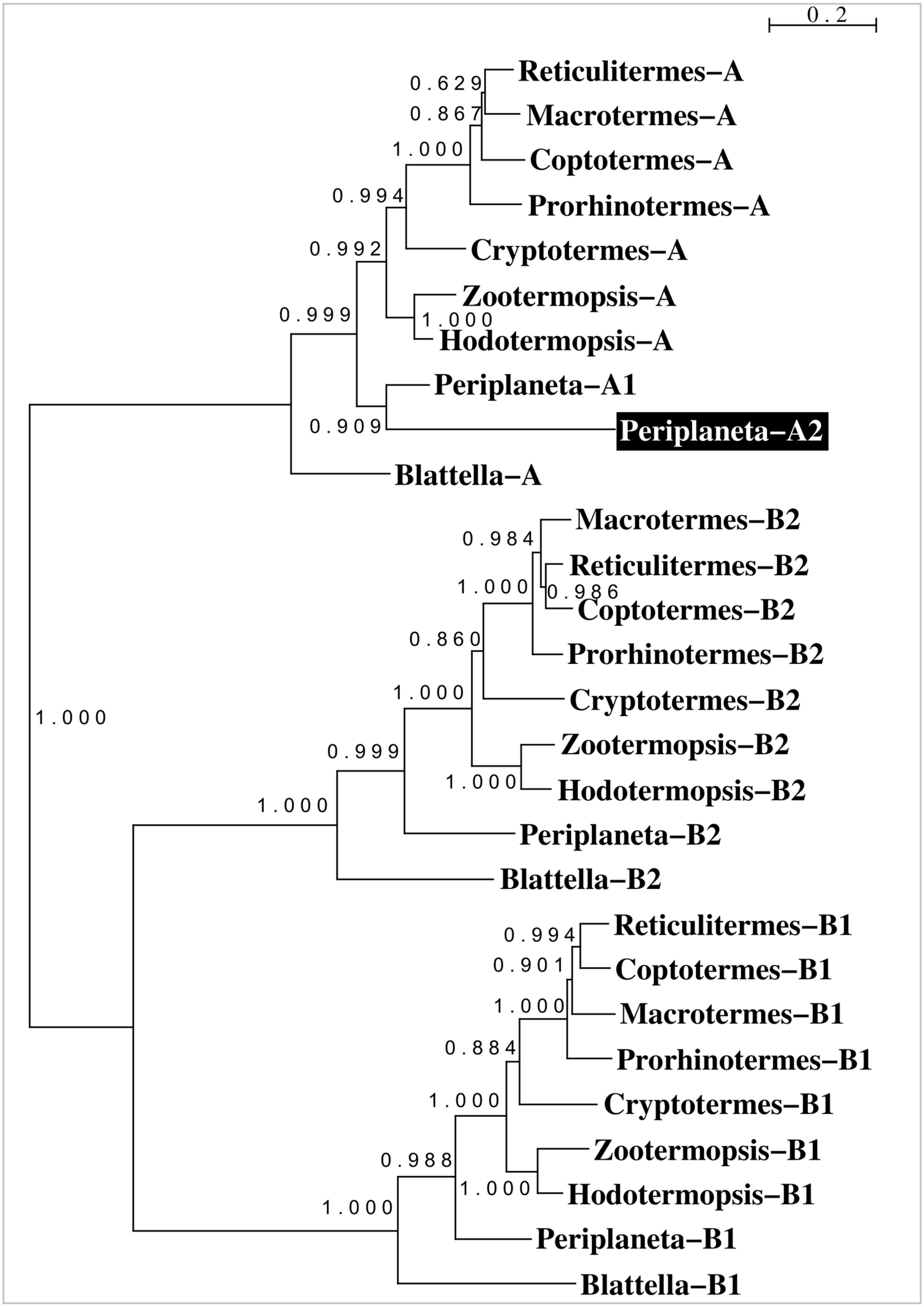


**Figure S12.** Phylogenetic tree of the termite insulin RTKs, showing that although *Periplaneta* has four such receptors, termites have only three.

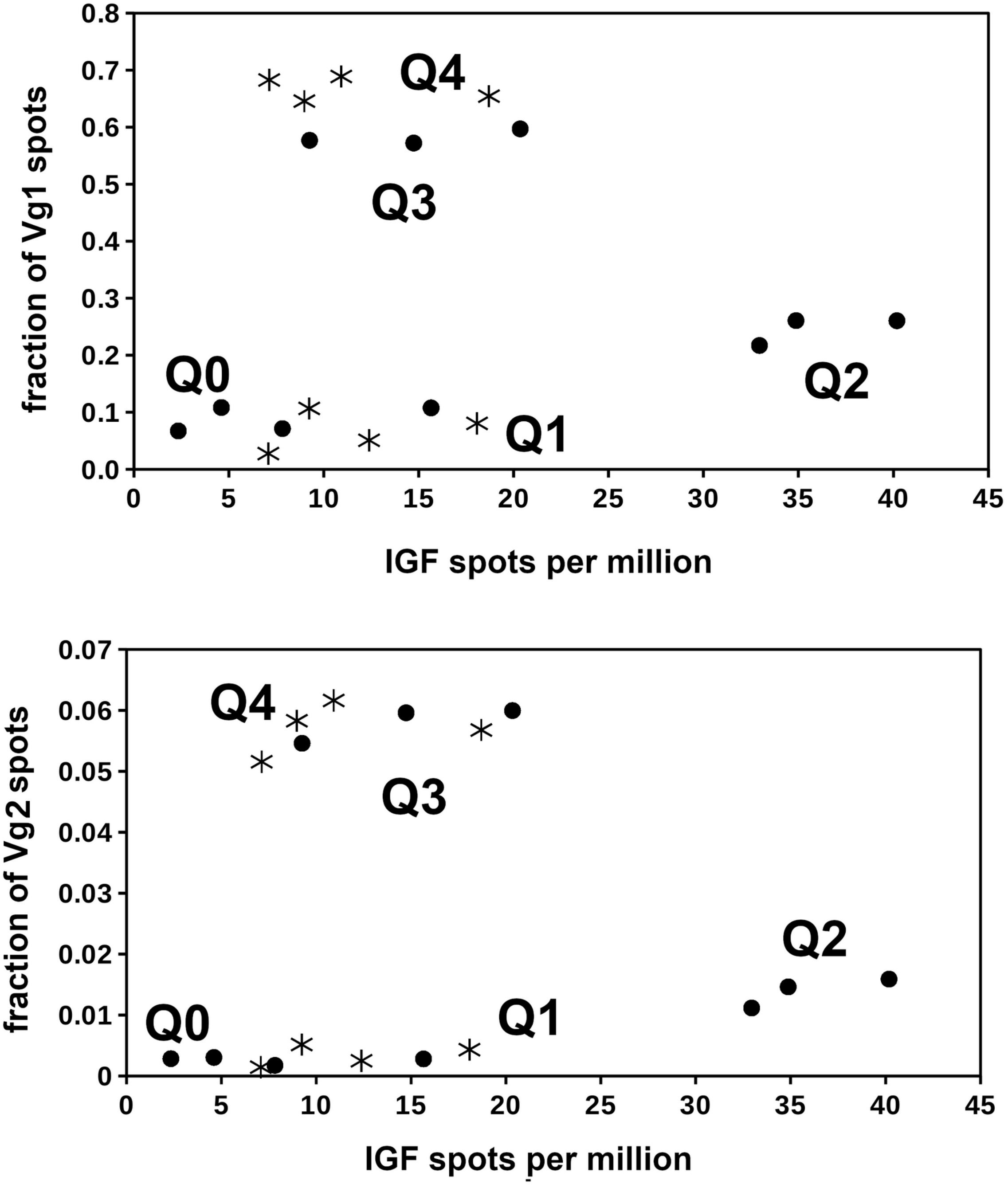


**Figure S13.** Top panel shows the correlation between the number of IGF and vitellogenin 1 spots in *M. natalensis* queens of different ages, Q0, Q2 and Q3 circles and Q1 and Q4 asterisks. Note that in Q0 and Q1 queens the fraction of vitellogenin 1 spots is similar, that it increases significantly in Q2 queens and then becomes very high in Q3 and Q4 queens. The bottom panel showa the correlation between IGF reads and vitellogenin 2, thus illustrating that qualitatively the results are identical for vitellogenins 1 and 2. Note that IGF expression appears highest in Q2 queens. The bottom panel of this figure was copied from figure 1 from Séité et al., 2022.

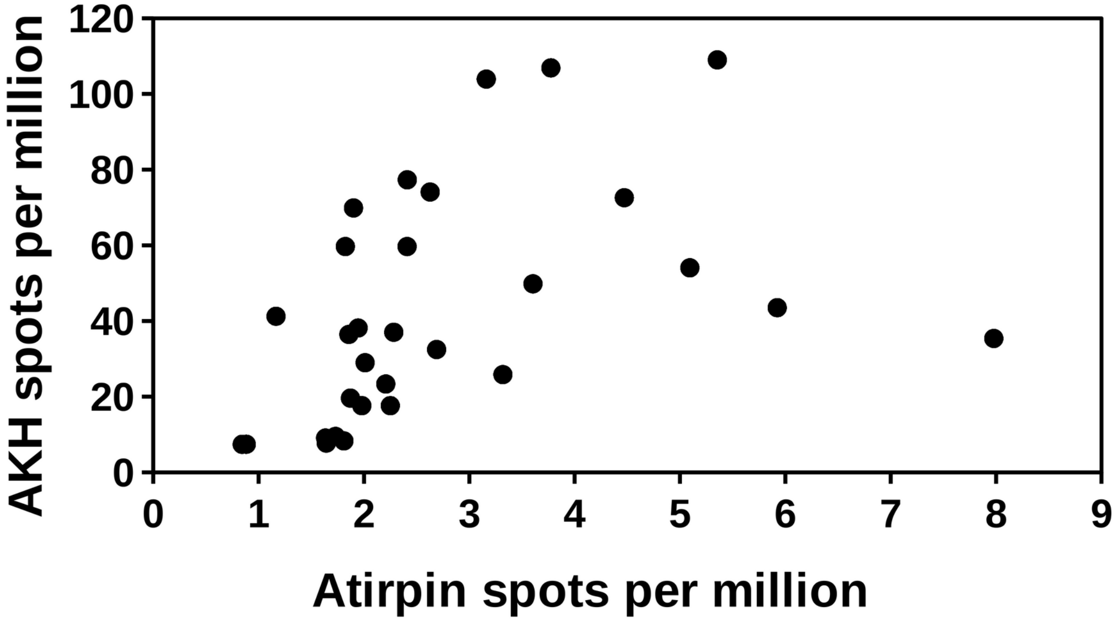


**Figure S14.** Data from *C. secundus* queens showing the correlation between spots for atirpin and AKH. Same data those in figs. 7-9 from the main text.
